# Direct reprogramming of the intestinal epithelium by parasitic helminths subverts type 2 immunity

**DOI:** 10.1101/2021.09.25.461778

**Authors:** Danielle Karo-Atar, Shaida Ouladan, Tanvi Javkar, Loick Joumier, Macy K. Matheson, Sydney Merritt, Susan Westfall, Annie Rochette, Maria E. Gentile, Ghislaine Fontes, Gregory J. Fonseca, Marc Parisien, Luda Diatchenko, Jakob von Moltke, Mohan Malleshaiah, Alex Gregorieff, Irah L. King

## Abstract

Enteric helminths form intimate physical connections with the intestinal epithelium, yet their ability to directly alter epithelial stem cell fate has not been resolved. Here we demonstrate that infection of mice with the symbiotic parasite *Heligmosomoides polygyrus bakeri* (*Hbp*), reprograms the intestinal epithelium into a fetal-like state marked by the emergence of *Clusterin*-expressing revival stem cells (revSCs). Organoid-based studies using parasite-derived excretory/secretory products reveal that *Hpb*-mediated revSC generation occurs independent of host-derived immune signals and inhibits type 2 cytokine-driven differentiation of secretory epithelial lineages that promote their expulsion. Reciprocally, type 2 cytokine signals limit revSC differentiation and, consequently, *Hpb* fitness indicating that helminths compete with their host for control of the intestinal stem cell niche to promote continuation of their life cycle.

## Main Text

Intestinal helminths remain pervasive throughout the animal kingdom by manipulating host defense pathways to prioritize tissue adaptation and repair over parasite expulsion (*1*). This host defense strategy begins at the gut epithelium: a monolayer of cells continually replenished by intestinal stem cells (ISCs) residing at the crypt base. As frontline effectors of barrier integrity, a key attribute of the ISC compartment is its ability to undergo extensive transcriptional reprogramming in response to injury (*2, 3*). Whether infectious organisms exploit ISC plasticity to persist in their hosts remains unknown. By interrogating *Heligmosomoides polygyrus bakeri* (*Hpb*) infection in mice, we demonstrate that these parasites directly reprogram ISCs, independent of host-derived factors. This reprogramming event coincides with adult parasite adherence to intestinal villi and is characterized by activation of a fetal-like, regenerative Hippo pathway signature and the emergence of *Clusterin*-expressing ‘revival’ stem cells (revSC). Morphological and single cell transcriptomic analyses of intestinal organoids exposed to *Hpb* excretory-secretory products (HES) revealed direct targeting of the ISC niche. HES also suppressed IL-13-induced secretory cell differentiation while deletion of type 2 cytokine signaling *in vivo* led to an epithelial-intrinsic expansion of revSCs and improved parasite fitness. Collectively, our study reveals how a helminth parasite co-opts a tissue development program to counter type 2 immune-mediated expulsion and maintain chronic infection.

A recent report demonstrated that expression of *Ly6a* (encoding Sca-1) marks epithelial cells undergoing a fetal-like reversion event in granuloma-associated crypts during the tissue-dwelling stage of *Hpb* infection (day 6 post-infection) (*4*). We performed a kinetic analysis of duodenal tissue from *Hpb*-infected adult C57BL/6 mice using RNAseq that confirmed this response, but revealed a more robust fetal gene signature during the luminal stage of infection, a time in which adult worms establish adherence to intestinal villi (day 14) (Fig 1A-B). To corroborate these results, we extracted intestinal crypts from *Hpb*-infected mice at different time points and cultured them under standard organoid growth conditions (Fig 1C). Adult intestinal organoids normally develop into polarized structures that mimic the crypt/villus architecture and are driven by abundant Lgr5+ ISCs (i.e. crypt-base columnar cells; CBCs) (*5*). Although crypts extracted at early time points post-infection yielded typical budding organoids with abundant Paneth cells (Fig 1D), crypts at day 14 post-infection developed into hollow spheres resembling fetal organoids (Fig 1D-E) (*6*). Furthermore, crypts and epithelial cells isolated at day 14 showed elevated expression of several fetal genes including *Clusterin* (*Clu*), *Il1rn* and *Msln* (Fig 1F, Fig S1A). These findings were confirmed using single molecule RNA *in situ* hybridization (RNAscope), which showed the loss of *Olfm4*+ ISC and a transient induction of *Ly6a* in granuloma-associated crypts at day 6, but sustained induction of *Il1rn* and *Il33* in distinct domains along the crypt-villus axis and the luminal stage-specific induction of *Clu* (Fig 1G-H, Fig S1B).

**Fig. 1.**
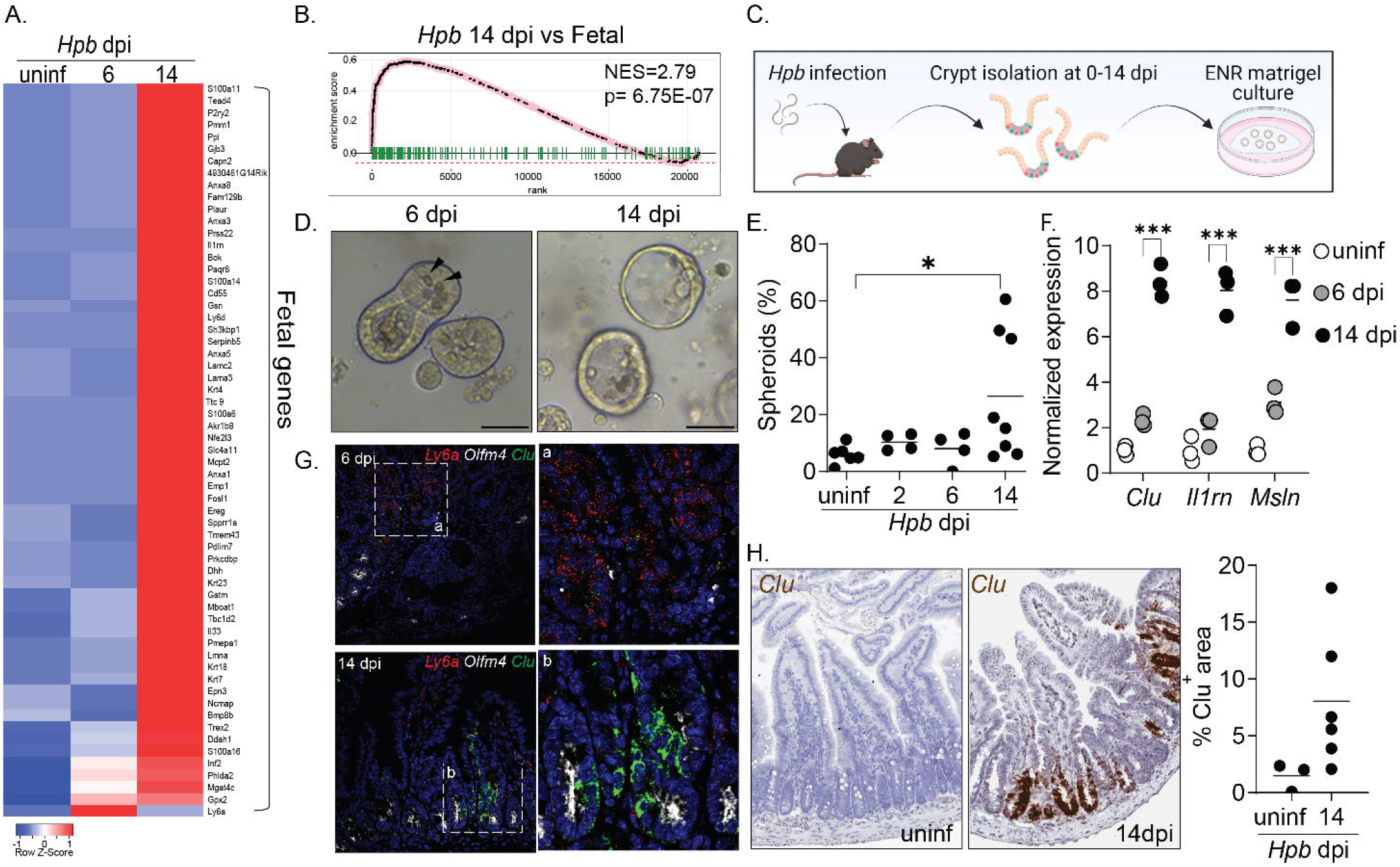
A fetal-associated transcriptional signature at the luminal stage of *Hpb* infection. (**A**) analysis of RNAseq dataset (*14*) showing reads per kilobase million (RPKM) of fetal-associated genes (*6*) in duodenal tissue at 6 and 14 days post-*Hpb* infection (dpi). Heat map was created using the Heatmapper web tool (*15*); uninf, uninfected. (**B**) GSEA of fetal-associated transcripts at 14 dpi. (**C**) C57BL/6 mice were infected with 150 L3 *Hpb* larvae, SI crypts were extracted at 2, 6 and 14 dpi and plated in Matrigel© domes for 24-48h; image created with Biorender.com. (**D**) representative photomicrographs of organoids cultured from *Hpb*-infected intestines, arrowheads indicate Paneth cells. (**E**) frequency of organoids with spheroid morphology. (**F**) qPCR analysis of organoids cultured from *Hpb* infected intestines. (**G**) RNAscope of fetal-associated transcripts in *Hpb*-infected tissue. a,b are enlarged insets from left panels. (**H**) colorimetric RNAscope of *Clu* mRNA and quantification of *Clu*+ areas. Scale bar, 50μm; N>3; *p<0.05, **p<0.01, ***p<0.005.

One interpretation of our findings is that luminal adult worms trigger fetal reprogramming because of the host immune response to parasite-induced breaches in the epithelium as previously suggested for the larval stages of infection (*4*). An alternative, but not mutually exclusive, possibility is that fetal reprogramming is driven by recognition of helminth-derived products by intestinal epithelial cells without *a priori* damage. To test the latter possibility, we stimulated organoids from the duodenum of uninfected mice with HES purified from adult *Hpb* parasites. Strikingly, HES-stimulated organoids robustly assumed a spheroid morphology with robust induction of *Clu* and other fetal markers, while also downregulating homeostatic markers of ISCs, *Lgr5* and *Olfm4* (Fig 2A-D). RNA sequencing followed by gene set enrichment analysis (fGSEA) confirmed the direct fetal-like reprogramming of organoids by HES (Fig 2E-F). Further analysis revealed that this reprogramming event was accompanied by the suppression of differentiated epithelial lineages (Fig 2G, Fig S1C-D) including Goblet and Paneth cells. Importantly, helminth-induced fetal-like reprogramming of the ISC niche was not observed in response to ES prepared from *Nippostrongylus brasiliensis* (NES), a rat parasite that briefly transits through the small intestine of infected mice (Fig 2H-J). Collectively, these results suggest that fetal reprogramming is associated with symbiotic parasites of the small intestine.

**Fig. 2.**
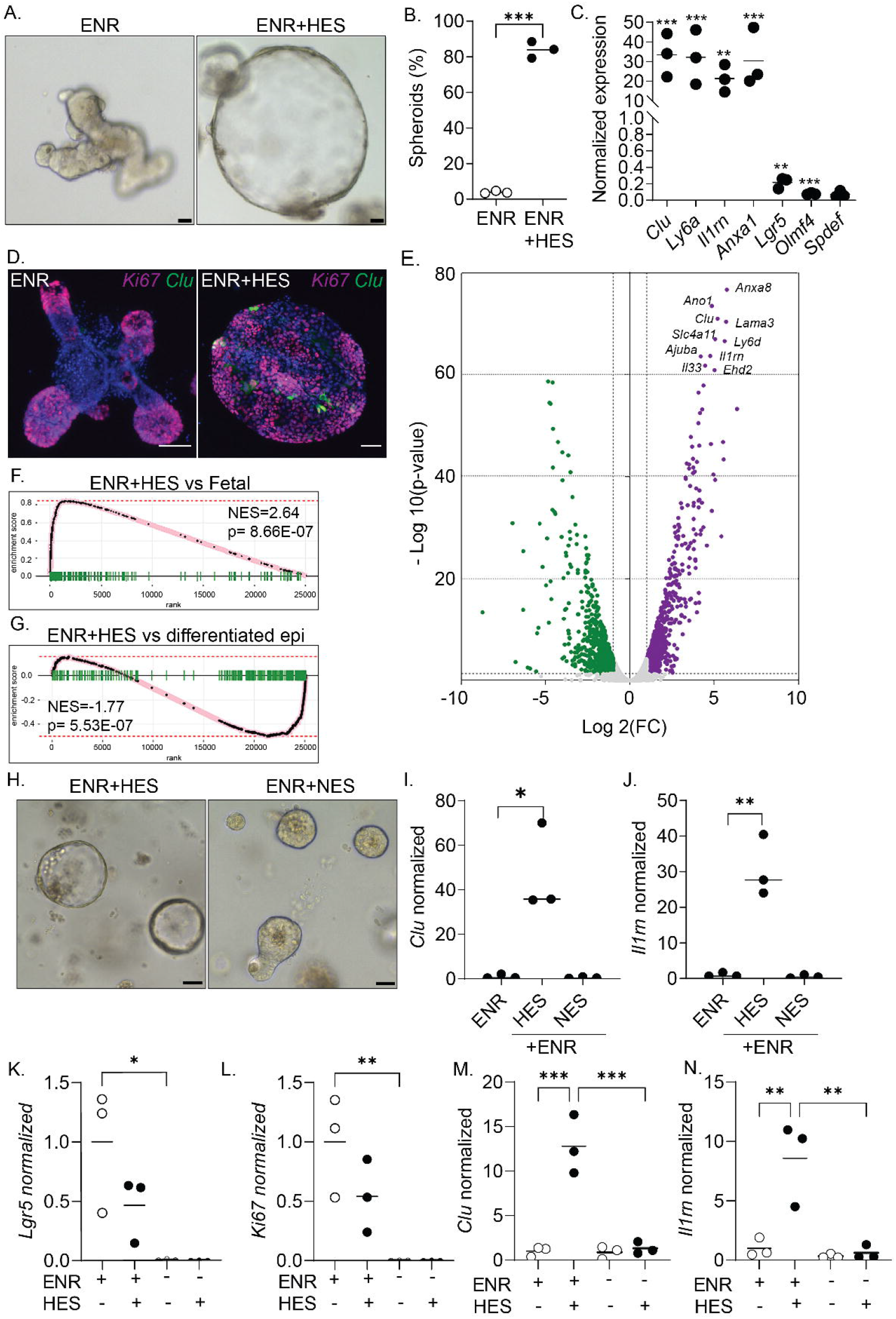
HES directly induces fetal reversion of intestinal organoids. (**A**) SI organoids were stimulated with ENR (control) or ENR+HES medium for 24h after three days in culture. (**B**) frequency of organoids with a spheroid morphology. (**C**) qPCR analysis of SI organoids stimulated with HES for 24h from the start of culture. (**D**) confocal photomicrographs of SI organoids from *Clu*^GFP^ crypts (*8*) stimulated with HES for 24h. (**E**) Volcano plot of differentially expressed genes from RNAseq analysis of HES-treated organoids, FDR<0.05. For a complete list of differentially expressed genes, see Table S1. GSEA of fetal-associated transcripts (**F**) and differentiated epithelial cell markers (**G**). Representative photomicrographs (**H**) and qPCR analysis (**I-J**) of SI organoids stimulated with NES for 24h. (**K-N**) qPCR analysis of SI organoids stimulated with HES for 24h in the presence or absence of ENR. Scale bar, 50μm; N=3; *p<0.05, **p<0.01, ***p<0.005.

Intestinal stem cells strictly depend on niche factors such as WNT and BMP morphogens, provided by crypt-adjacent stroma and Paneth cells, to regulate their self-renewal and lineage commitment (*2*). Therefore, EGF, Noggin and R-spondin-1 (ENR) are an integral part of the organoid culture medium. To assess whether organoids depend on niche factors for helminth driven reprogramming, organoids grown with ENR were re-plated in the absence of niche factors and stimulated with HES. While the removal of niche factors resulted in an expected decrease in proliferation and loss of the homeostatic stem cell marker *Lgr5*, HES was unable to initiate a fetal transcriptional response (Fig 2K-N). Thus, HES-mediated reprogramming requires the presence of niche factors and likely targets a mitotic progenitor cell.

The Hippo signaling pathway is a well-established regulator of gut organogenesis and tissue regeneration in response to damage and has been linked to fetal reprogramming (*7*). Consistently, *Hpb* induced nuclear translocation of YAP, a transcriptional effector of Hippo signaling, in the intestinal epithelium at the luminal stage of infection (Fig S2A). While many of the fetal-associated genes including *Clu* are YAP target genes, HES-induced fetal-like reprogramming of organoids and *in vivo Clu* induction were only partially *Yap*-dependent (Fig S2B-D and data not shown). Furthermore, treatment of organoids with recombinant CLU was not sufficient to induce spheroids and stimulation of *Clu*-deficient organoids with HES did not prevent organoid reprogramming (Fig S2E-G and data not shown). Thus, *Clu* expression represents a broader transcriptional network mediating fetal-like reprogramming that does not exclusively rely on Yap-dependent signaling.

For a more in-depth analysis, we performed single-cell RNA sequencing (scRNAseq) on control and HES-stimulated organoids. Our analysis yielded eight cell clusters, identifying stem (*Lgr5*+ CBCs) and progenitor cells as well as mature secretory populations (Fig 3A-B, Fig S3A-B). In accordance with our bulk RNAseq results (Fig 2), HES expanded a distinct population enriched in fetal genes, which was rare in control treated organoids and that we previously identified as revival stem cells (revSCs) in irradiated crypts (*8*). By contrast, Goblet and Paneth cell clusters were drastically reduced in HES-stimulated organoids while secretory progenitor cells (marked by *Atoh1* and *Spdef* expression) were enriched (Fig 3A, C). Trajectory analyses established the displacement of Transit Amplifying (TA) cells and emergence of an altered and shorter path towards revSCs in HES-stimulated organoids (Fig 3D). To validate the importance of revSCs in driving renewal of the gut epithelium upon *Hpb* infection *in vivo*, we performed lineage-tracing studies using tamoxifen-inducible *Clu* fate-mapping mice (*8*). As shown in Fig 3E-G, *Clu*+ revSCs were activated specifically at the luminal stage of infection giving rise to Tomato+ ribbons comprising the entire crypt-villus axis, while minimal tracing events were detected at earlier time points. These data suggest that *Hpb* parasites target CBC and TA cells to support revSC generation while blocking the generation of secretory cell lineages with direct anti-helminth activity.

**Fig. 3.**
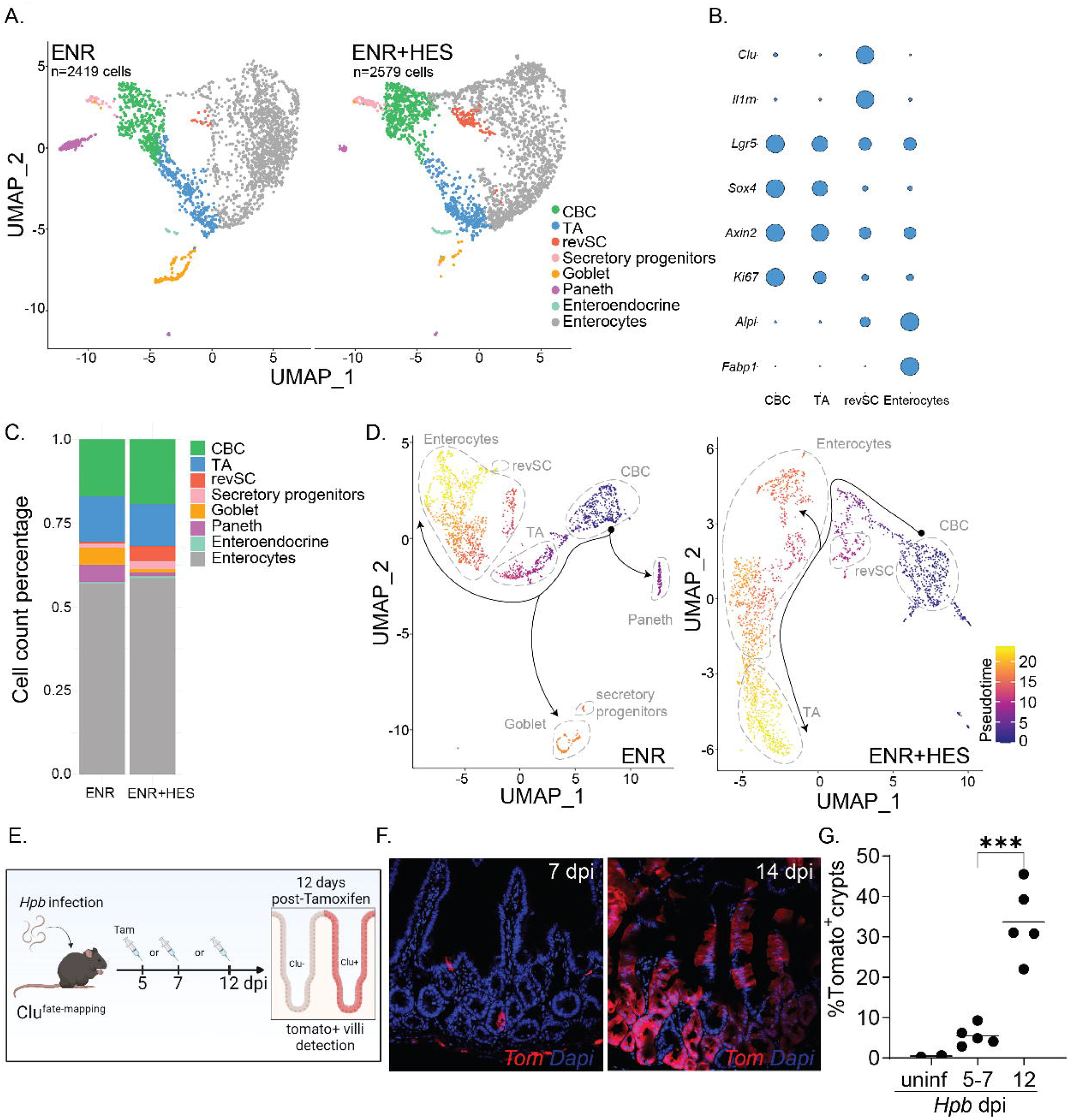
HES expands revSCs and limits secretory cell differentiation. SI organoids were stimulated with HES for 24h from the start of culture and scRNAseq analysis was performed. (**A**) UMAP projection plots; colors represent cells clustered together based on gene expression similarity. (**B**) Representative cluster identifying markers; circle size represents average expression of indicated transcripts (complete plot and gene expression list can be found in Fig S3B and Table S2). (**C**) the proportion of each cluster within each sample is presented. (**D**) trajectory analysis using CBCs as the pseudotime source variable (pseudotime = 0). (**E-G**) *Clu* fate-mapping (*Clu*-CreERT; Rosa26-LSL-tdTomato) (*8*) mice were infected with *Hpb* and Clu+ dtTomato-expressing cells (Tom) were induced by Tamoxifen injection at 5, 7 or 12 dpi and imaged at day 12 post-Tamoxifen. N=2-5; *p<0.05, *p<0.01, ***p<0.005.

Tuft and Goblet cells are secretory epithelial lineages that orchestrate the initiation and effector functions, respectively, of the type 2 immune response that is responsible for parasitic helminth expulsion (*9*–*11*). The main driver of secretory cell differentiation is STAT6-dependent type 2 cytokine signaling (via IL-4 and IL-13) (*12*). Building on our scRNA sequencing results, we assessed whether HES was able to directly inhibit type 2 cytokine-induced differentiation of secretory cells. Compared to IL-13 stimulation of organoids which resulted in robust *Spdef, Clca1* (encoding Gob5*), Muc2* and *Dclk1* mRNA expression, the addition of HES prevented induction of these lineage-specific transcripts (Fig 4A-D). These results were confirmed by immunofluorescence microscopy (Fig 4G). However, when organoids were pre-treated with IL-13 to promote secretory cell differentiation, subsequent exposure to HES was unable to induce fetal gene expression (Fig 4E-F). Therefore, HES and type 2 cytokines act in a counter-regulatory manner on the ISC niche.

**Fig. 4.**
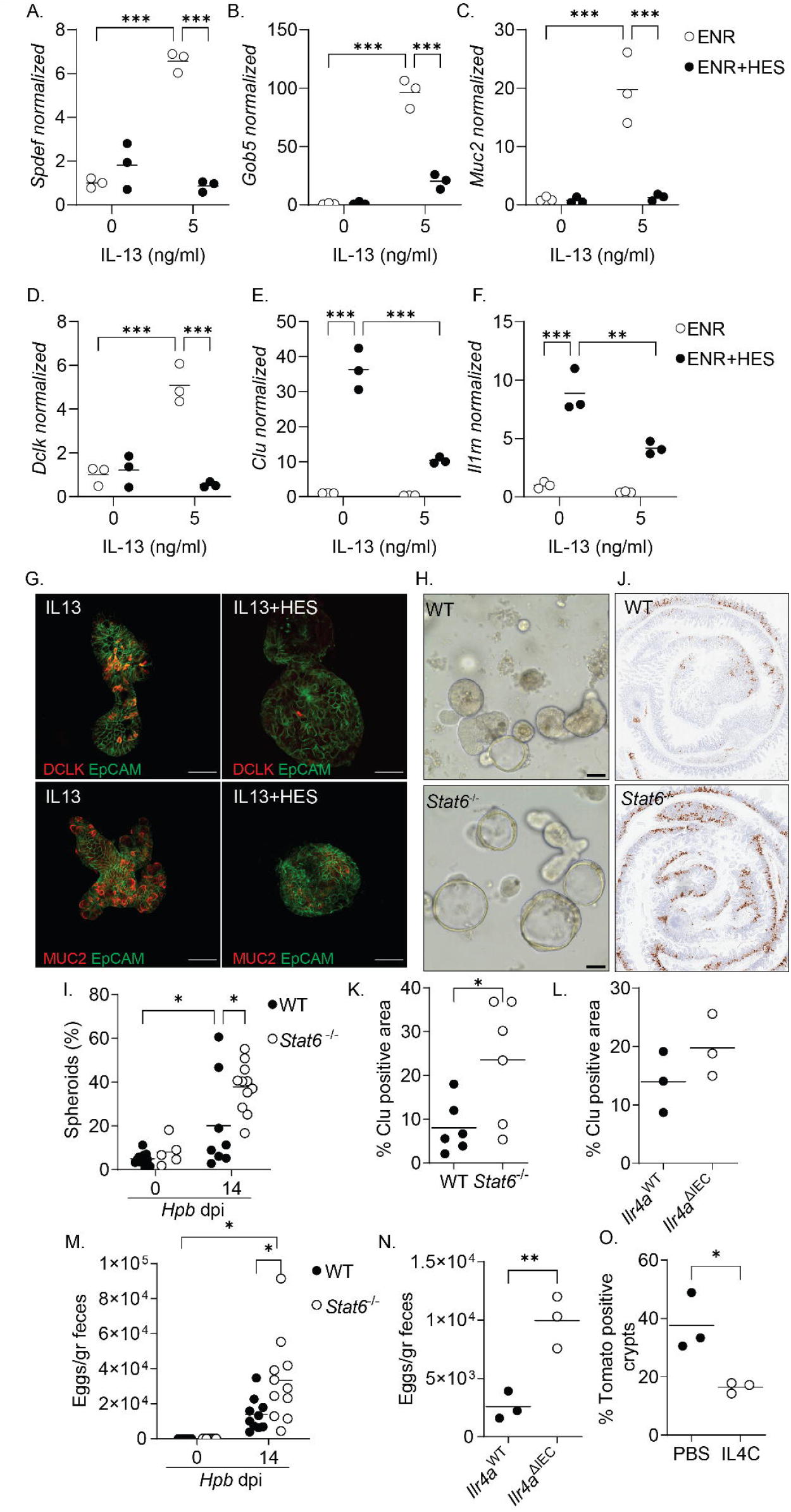
Counter-regulation of the ISC niche by helminths and type 2 cytokine signaling. (**A-D**) SI organoids were stimulated at the start of culture with HES +/- IL13, qPCR analysis was performed at 48h. (**E-F**) SI organoids were stimulated with IL13 for 10 days prior to HES stimulation and qPCR analysis was performed. (**G**) photomicrographs of organoids stimulated with HES +/- IL13 for 48h. (**H-I**) C57BL/6 and *Stat6*^-/-^ mice were infected with *Hpb* and crypts were extracted at 14 dpi and plated in Matrigel© domes for 24h. Representative photomicrographs (**J**) and quantification (**K**) of *Clu*+ areas of WT and *Stat6*^-/-^ tissues at 14 dpi. (**L**) Quantification of *Clu*+ areas of *Il4ra*^WT^ and *Il4ra*^ΔIEC^. (**M-N**) Fecal *Hpb* egg counts from *Stat6*^-/-^ and *Il4ra*^ΔIEC^ mice. (**O**) Quantification of *Clu*+ areas of IL4C-treated mice. Scale bar, 50μm; N>3; *p<0.05, **p<0.01, ***p<0.005.

To test the involvement of type 2 immune signaling in the regulation of *Hpb*-induced fetal-like reprogramming *in vivo*, we infected *Stat6*-deficient mice with *Hpb*. Remarkably, loss of STAT6 resulted in increased *ex vivo* fetal-associated spheroid formation and *in situ* intestinal *Clu* expression (Fig 4H-K). To specifically test the role of epithelial-intrinsic type 2 immune signaling in the induction of *Clu*+ revSCs, Villin^Cre^*Il4ra*^fl/fl^ mice (*Il4ra*^ΔIEC^) were infected with *Hpb* and a trend towards increased *in situ Clu* expression was detected (Fig 4L, Fig S4A). We next assessed *Hpb* worm fitness and persistence by evaluating adult *Hpb* burden in *Stat6*^-/-^ mice (Fig S4C) and fecal egg counts in *Stat6*^-/-^ and *Il4ra*^ΔIEC^ mice (Fig 4M-N). In association with enhanced *Clu* expression, *Stat6*^-/-^ and *Il4ra*^ΔIEC^ mice had elevated worm and egg burdens at day 14 post-infection suggesting that epithelial-intrinsic type 2 immune signaling is needed to limit the induction of revSCs and worm persistence. To control for differences in parasite fitness, we took a gain of function approach and treated Clu-fate mapping mice with low dose IL-4 complexes (IL4C) during *Hpb* infection. This transient amplification of type 2 cytokine signals led to a reduction of *Hpb*-induced Tomato+ ribbons without a difference in *Hpb* egg production (Fig 4O, Fig S4B,D). Finally, to assess the immune landscape in *Stat6*^-/-^ and *Il4ra*^ΔIEC^ mice following *Hpb* infection, we performed immune-phenotyping of the small intestine (SI), Peyer’s patches (PPs) and mesenteric lymph nodes (MLN) by flow cytometry (Fig S5). As expected, accumulation of GATA3+ Th2 cells in the MLN, PPs and SI of *Hpb*-infected animals was largely eliminated in *Stat6*^-/-^ mice. By contrast, infected *Il4ra*^ΔIEC^ mice showed similar numbers of Th2 cells to littermate controls indicating that direct type 2 cytokine signaling, not immune deviation *per se*, is responsible for driving epithelial reprogramming. Taken together, our data establish a balanced regulation of the ISC niche by helminths and type 2 immune signaling that supports durable infection.

Expulsion of multicellular helminths from the intestinal lumen requires production of type 2 cytokines that direct the epithelium to increase mucus-producing goblet cells and enhance the rate of epithelial turnover, the so-called “weep and sweep” response (*13*). It is well-documented that helminths secrete immunomodulatory factors that target type 2 cytokine-producing cells to limit anti-helminth immunity (*1*). Here we demonstrate that helminths also evade expulsion through direct reprogramming of the intestinal epithelium into a fetal-like state. Although fetal reversion of the intestine is critical for regeneration upon injury, we propose that symbiotic parasites such as *Hpb* that have evolved with their hosts, co-opt this restorative pathway to persist and continue their life cycle (Fig S6). Our study provides a new conceptual framework that may not only lead to new anthelmintics that interfere with parasite-epithelium signaling networks, but also spur helminth-based therapies that aim to rejuvenate the intestinal barrier following acute injury.

## Acknowledgments

We thank all members of the King and Gregorieff laboratories for their support and feedback in preparing this manuscript. We greatly appreciate the technical support of the MUHC-RI Animal Resource Division and the Histology platform.

## Author contributions

Conceptualization: ILK, AG, DKA

Methodology: ILK, AG, DKA, MM, LD

Investigation: DKA, TJ, SO, SM, GF, AR, LJ, MP, SW, MG, MKM, GJF

Funding acquisition: ILK, AG

Writing – original draft: ILK, AG, DKA

Writing – review & editing: All authors

## Data and materials availability

Data sets present will be uploaded at a later time.

## Supplementary Materials

Materials and Methods

Supplementary Text

Figs. S1 to S6

Tables S1 to S2

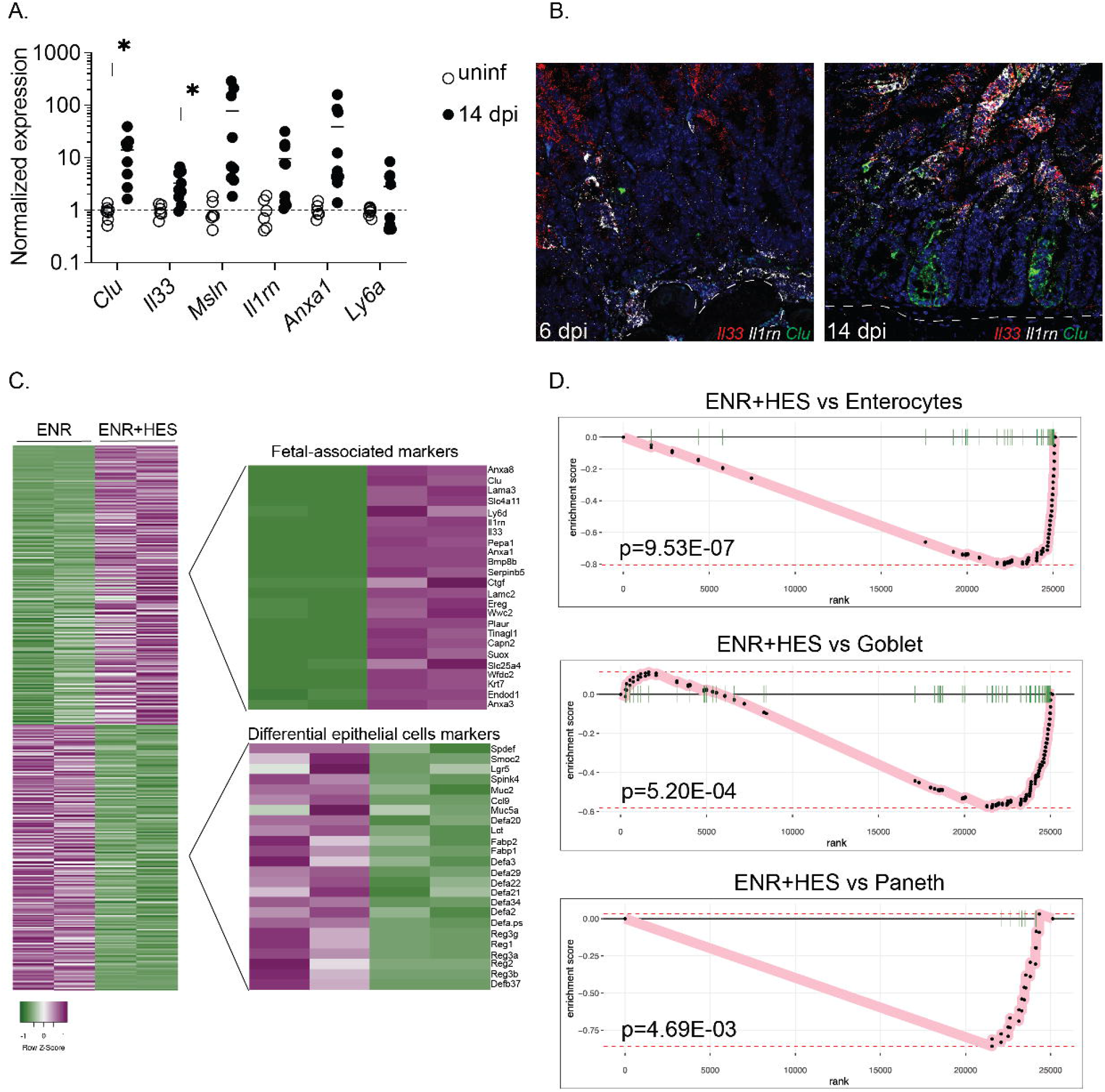

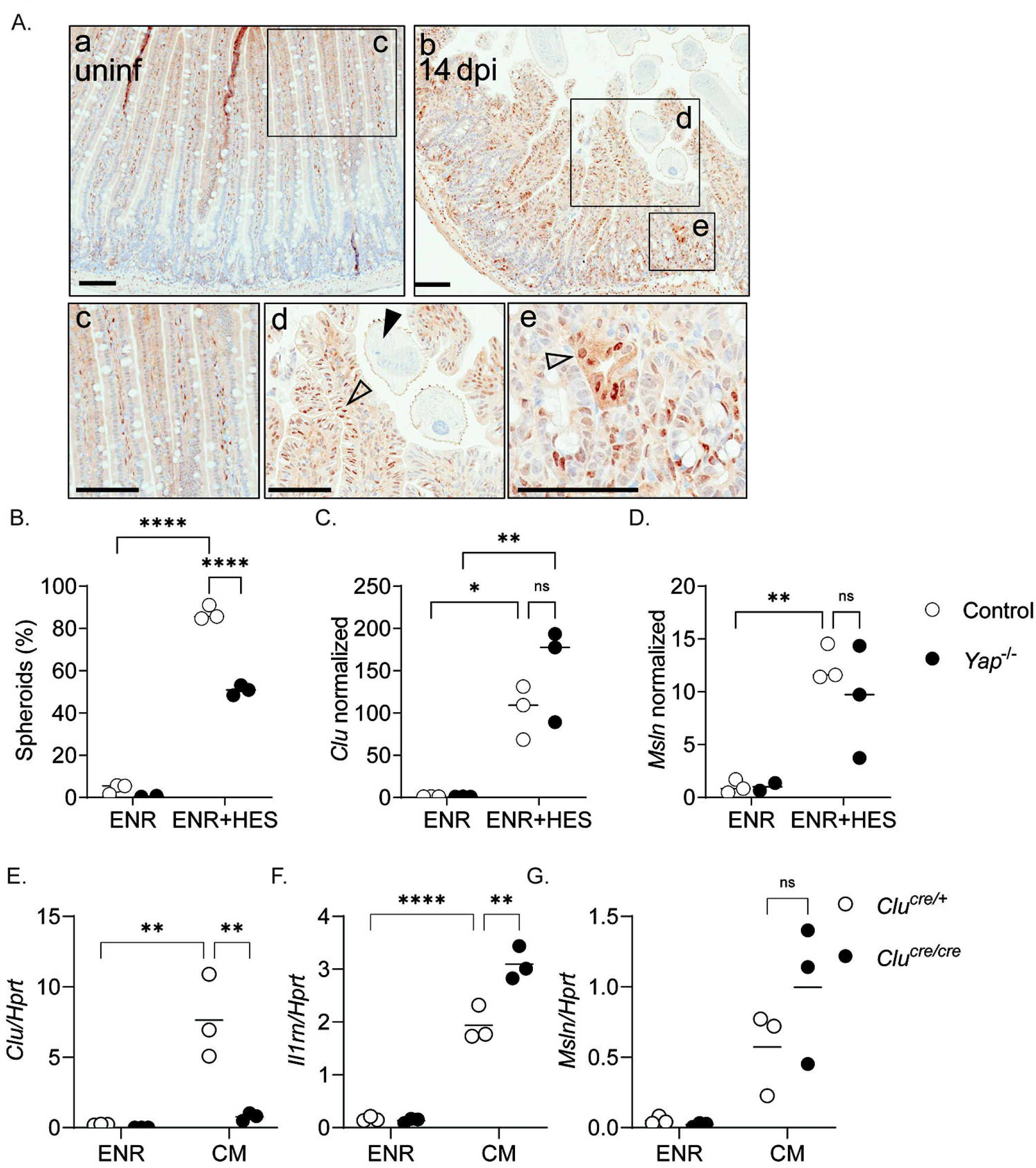

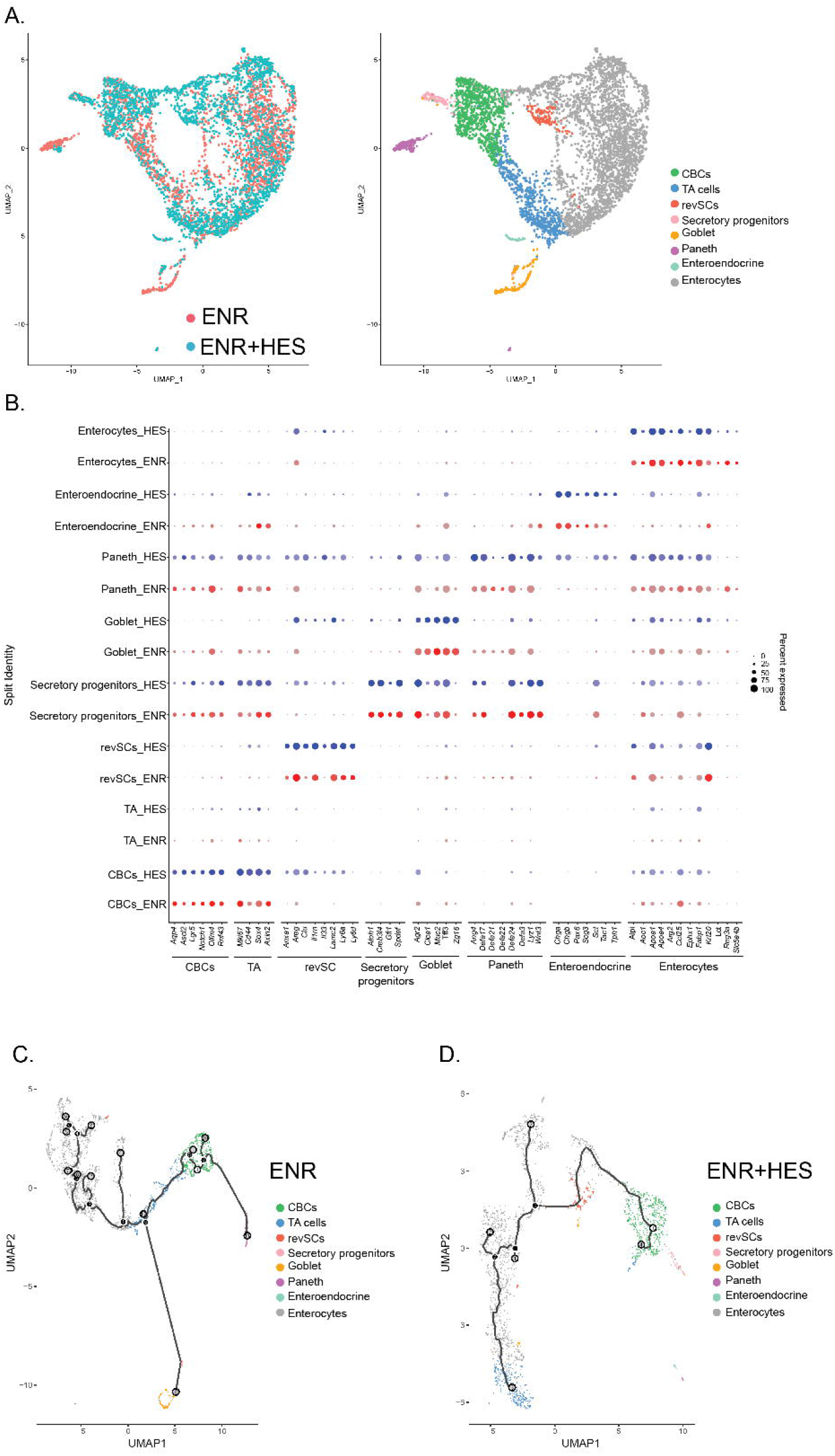

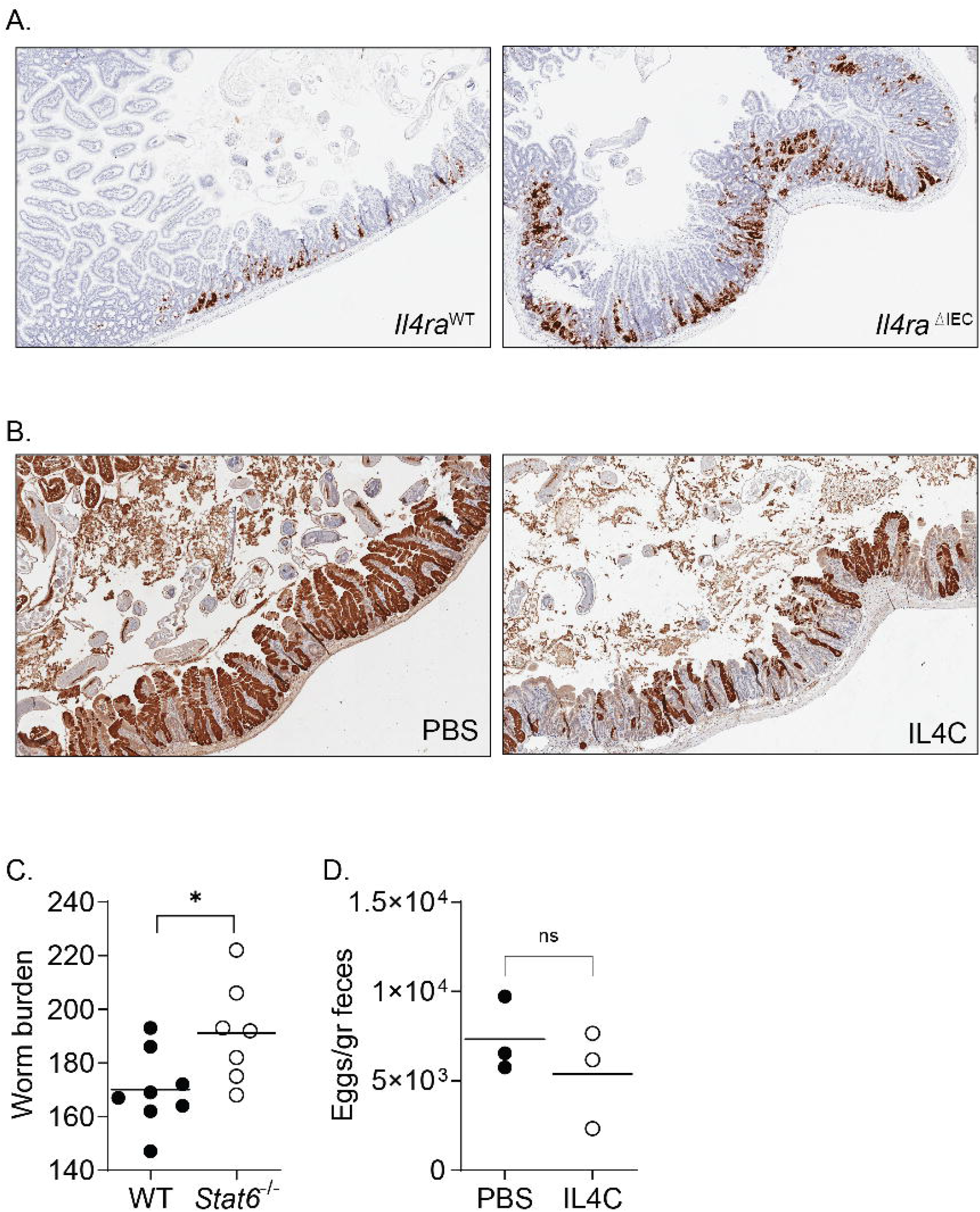

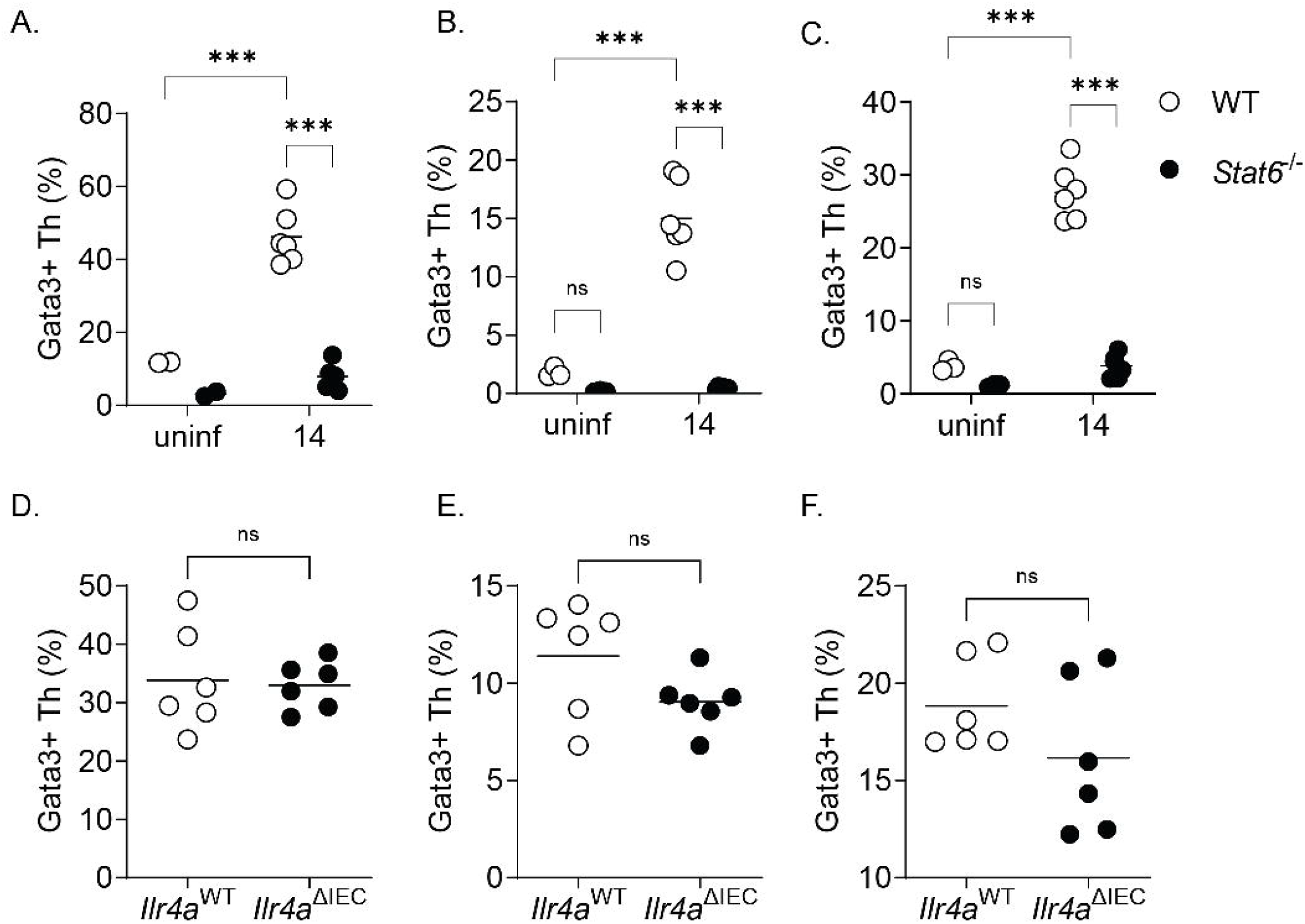

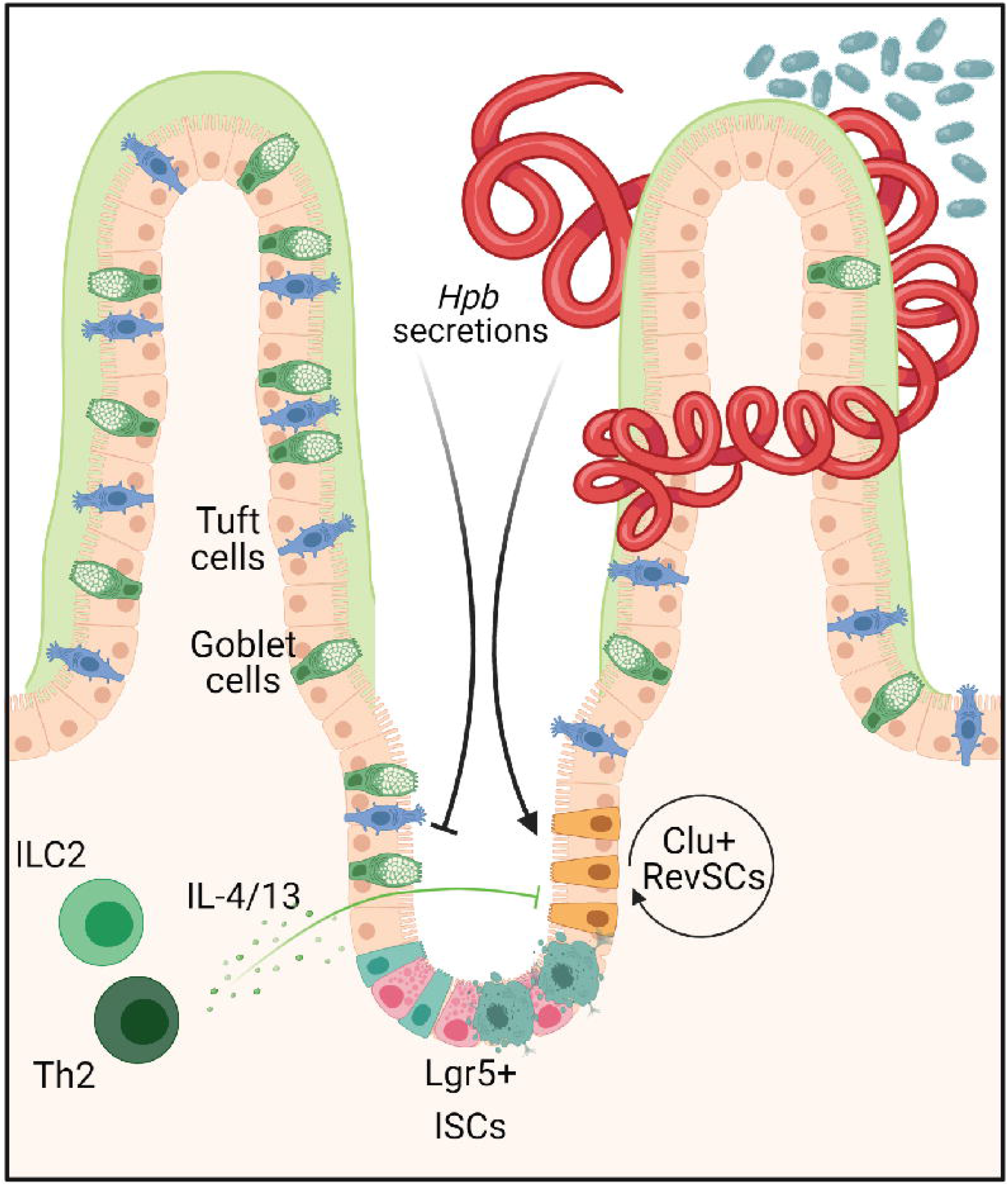

