## Supplementary material for "Direct reprogramming of the intestinal epithelium by parasitic helminths subverts type 2 immunity": Methods and supplementary material

### Materials and Methods

Animals

All experiments were performed in accordance with the McGill University Health Center Animal Resource Division. Wild type (WT) C57BL/6, Stat6 $^{-/-}$ and Yap$^{dupa/dupa}$ mice were obtained from the Jackson Laboratory. *Villin*$^{CRE}$ mice were kindly provided by Nicole Beauchemin (McGill University). *Il4ra*$^{fl/fl}$ mice were kindly provided by Frank Brombacher (International Center for Genetic Engineering and Biotechnology, Cape Town). *Villin*$^{CRE}$*Il4ra*$^{fl/fl}$ were bred in house. *Clu*-CreERT; Rosa26-LSL-tdTomato, *Clu*$^{GFP}$ and *Clu*$^{cre/cre}$were generated as previously described (*8*). All mice were bred and maintained under specific pathogen-free conditions. Female and male mice of 6–10 weeks of age were used.

Helminth infection

For *Heligmosomoides polygyrus bakeri* (*Hbp*) infection, mice were infected by gavage with 150 L3 stage larvae diluted in sterile water, killed at the indicated time points, and tissues were harvested for analysis. For histology the proximal 5 cm of the small intestine were harvested after the indicated days, flushed, and fixed in 10% formalin for 24 hours before standard FFPE processing. For *Nippostrongylus brasiliensis* infection, infectious third-stage larvae (L3) were maintained by passaging through Lewis rats (Envigo), as previously described (*16*). For histology of *N. brasiliensis*-infected intestines, mice were infected subcutaneously with 500 L3 larvae. The proximal 10-12 cm of the small intestine were harvested after 7 days, flushed, and fixed in 10% formalin for 2 hours. Intestines were then rolled into "swiss rolls", placed in a tissue cassette, and fixed for an additional 18-24 hours before standard FFPE processing.

Helminth excretory-secretory products preparation

To generate HES wild type mice were infected by gavage with 400 L3 *Hpb* larvae, and adult worms were manually harvested from the small intestine after 21 days as previously described (*17*). Sterile adult worms were than cultured overnight in organoid basal medium at approximately 130 worms/ml. The following day, culture medium containing HES was removed, filtered through a 0.2μm filter and kept at minus 80$^{0}$C until use. For NES preparation, rats were infected subcutaneously with 3000 L3 *N. brasiliensis*, adult worms were manually harvested from the small intestine after 7 days and cultured as described for HES.

Small intestinal organoids culture

The small intestine (SI) was harvested, flushed with PBS, cut into 3 approximately 10cm pieces, and opened longitudinally. Intestine pieces were rinsed with PBS, and then incubated in 2mM EDTA at 4 °C for 30 min with gentle agitation. Crypts were further released from the SI by vigorous shaking in 10mL PBS, three times, and passed through a 70μm cell strainer. Isolated crypts were resuspended in 10mL basal organoid medium (Advanced DMEM/F12 (Gibco), 10 mM HEPES (Gibco), 1× GlutaMAX (Gibco), 1% penicillin/streptomycin (Gibco). Crypts were plated in 30μL Matrigel (Corning) domes in 24-well plates and 500μL of culture medium was added to each well. Organoids culture medium (ENR): basal organoid medium with the addition of N2 Supplement (1:100, Gibco), B27 Supplement (1:50, Gibco), 1mM N-Acetylcysteine, 50ng/mL recombinant

Epithelial Growth Factor (EGF), in-house made R-spondin 1 and Noggin conditioned mediums (1:50). Organoids were maintained for the indicated times and passaged every 3-4 days by physical dissociation of the Matrigel and replating in fresh ENR medium. For HES and/or NES stimulation, HES and/or NES were supplemented with N2 (1:100, Gibco), B27 (1:50, Gibco), 1mM N-Acetylcysteine, 50ng/mL recombinant Epithelial Growth Factor (EGF), in-house made R-spondin 1 and Noggin conditioned mediums (1:50). For IL13 stimulation, organoids were treated with 5ng/ml of IL13 simultaneously with HES for 24h or pre-treated with 5ng/ml IL13 for 10 days following the addition of HES post-splitting for 18h.

#### ENR withdrawal

Established organoids were deprived of growth factors (R-spondin1, Noggin and EGF) post-splitting for 24 hours. Organoids were grown either in complete ENR media, complete ENR media with HES, deprived media or deprived media with HES for 18 hours.

#### Organoids whole-mounting and confocal microscopy

Established organoids were passaged and replated in a 1:1 ENR:Matrigel solution; 30µL domes were plated on Nunc™ Lab-Tek™ II Chamber Slide™ and incubated overnight in 200µL media at 37°C. Domes were fixed in 10% Formalin for 30 minutes at room temperature (RT), and permeabilized with PBS-T (PBS + Triton 0.5%) for 15 minutes. The samples were blocked with 200µL blocking solution (3% BSA in PBS) for 1h at RT. Organoids were incubated with primary antibodies overnight at 4 °C, followed by incubation with secondary antibodies at RT for 1 h. The nuclei were stained with 4′,6-diamidino-2-phenylindole (DAPI; 1ug/ml). All fluorescent images were taken at 20X after mounting using a Zeiss Confocal LSM700. Specific antibodies include: Anti-Mouse/Rat Ki67 eFluor 660 (Invitrogen), Anti-mouse CD326 (EpCAM) Alexa Fluor 488 (BioLegend), Goat anti-GFP (Invitrogen), Anti-goat IgG Alexa Fluor 555 (Invitrogen), Rabbit anti mouse Dclk (Abcam), Rabbit anti mouse Muc2 (Abcam) and Goat anti Rabbit IgG Alex Fluor 555.

#### RNAscope

RNAscope was performed according to manufacturer instructions (ACD bio).
RNAscope probes: *Clu*: cat# 427891; *Il1rn*: cat# 495101; *Il33*: cat# 400591-C2; *Ly6a*: cat# 427571-C2. Quantification of Clu positivity by colorimetric RNAscope was performed using the pixel count function of Qupath, a digital pathology image analysis software (*18*).

#### RNA extraction and qPCR

RNA was extracted using Tri reagent (Sigma) or PureLink™ RNA Mini kit (Invitrogen) according to manufacturer instructions.
Primers used:
Clu Fw: aaaccaacgcagagcgcaag
Clu Rv: ctccagagcatcctctttcttctt
Il1rn Fw: aggcatgtgctctaccatcatgct
Il1rn Rv: gccgacatggaataaggctggc
Msln Fw: tctccaaacagtggtgtggg
Msln Rv: aagcagtaggaagcttcggc
Lgr5 Fw: agagcctgataccatctgcaaac

Lgr5 Rv: tgaaggtcgtccacactgttgc
Olfm4 Fw: gctcctggaagctgtagtca
Olfm4 Rv: ggccccaggcaccatattta
Gob5 Fw: catcgccatagaccacgacg
Gob5 Rv: ttccagctctcgggaatcaaa
Muc2 Fw: ctgaccaagagcgaacacaa
Muc2 Rv: catgactggaagcaactgga
Spdef Fw: gctcctggaagctgtagtca
Spdef Rv: ggccccaggcaccatattta
Dclk Fw: aacgtcaagaccacctcagc
Dclk Rv: gagccgtcttcttgttcagc
Il33 Fw: gacacattgagcatccaagg
Il33 Rv: tgattgacttgcaggacagg
Ly6a Fw: ggaggcagcagttattgtgg
Ly6a Rv: gctacattgcagaggtcttcc
Anxa1 Fw: caaccatcgtgaagtgtgcc
Anxa1 Rv: atgccttatggcgagttccg
Ki67 Fw: cctggtcaccatcaagcgg
Ki67 Rv: attcaatactccttccaaacaggc

Bulk RNA sequencing

SI organoids were stimulated with ENR (control) or ENR+HES medium for 24h at day 0 of culture. RNA was extracted using Tri reagent (Sigma) and quality was checked with the Bioanalyzer RNA 6000 kit. Libraries were prepared using the NEB mRNA stranded library prep Kit (New England Biolabs) and paired-end sequenced on NovaSeq 6000 at 25M reads per sample. RNA sequencing reads were aligned to the mm10 reference genome using STAR (*19*). Read counts were calculated using the strand specific exonic reads of each gene and duplicate genes were merged using HOMER (*20*). Transcripts per million (TPM) was used to evaluate the correlation among replicates for quality control. Differential gene expression was calculated from the raw read counts using edgeR (*21*).

Gene set enrichment analysis

For pathway analyses, custom gene sets from genes differentially (up- or down-regulated) expressed in previous cell-type specific transcriptomic studies were build. Fetal-associated gene list was built using Table S1 from Mustata et al (*6*). Differentiated epithelial cell marker list was built using Table S4 from Haber et al (*22*). Transcriptomes from this work were analyzed for enrichment in differentially expressed genes in the above mentioned gene sets using the 'fgsea' R package (https://github.com/ctlab/fgsea) (*23*).

Single cell RNA sequencing

SI organoids were stimulated with ENR (control) or ENR+HES medium for 24h at day 0 of culture. Single cell suspensions were prepared by manual disturbance of Matrigel domes followed by incubation in TrypLE (Gibco) for 40 minutes at 37$^0$C, were loaded on a Next GEM Chip G (PN-1000120) together with Next GEM Single Cell 3′ GEM Kit v3.1 (PN-1000121) and single cells were captured on a 10X Genomics Chromium controller with a recovery target of 5000 cells per sample. cDNAs were generated following the 10X

Genomics protocol. cDNAs were size selected using SPRIselect beads from Beckman Coulter (B23318), and their quality was checked with a Bioanalyzer High Sensitivity DNA Kit from Agilent (5067-4626). One quarter of the total cDNA was used to generate libraries using Next GEM Single Cell Library Kit (PN-1000121) and barcoded using the Single Index Kit Set A (PN-111213) following the 10X protocol. Libraries were size selected using SPRIselect beads and their quality was checked with the Bioanalyzer High Sensitivity DNA Kit. Libraries size were centered at 440 bp, and paired-end sequenced on NovaSeq 6000 at 125M reads per sample. Single-cell matrices were generated using Cellranger (v.4.0.0) provided by 10X genomics. Sequences were aligned with the mm10 mouse transcriptome. Matrices were imported and analyzed with the R package "Seurat" (v.4.0.3) (*24*). We filtered out genes expressed in less than 3 cells and low-quality cells expressing less than 200 genes. Cells with more than 100000 transcripts were removed as they likely represent cell doublets (*24*). In addition, immune cells (marked by Cd3g) were removed from the analysis. Each matrix was normalized with Seurat's "NormalizeData" function using "LogNormalize" as the normalization method and a scale factor equal to 10000 (default value). Matrices were integrated using the "IntegrateData" function. Linear dimensionality reduction was performed using the "RunPCA" function. Non-linear dimensionality reduction was then performed using the "RunUMAP" function, considering the Euclidean distances of the first 30 PCA components. kNN graph construction was performed using the "FindNeighbors" function, which also considered the first 30 PCA components. Clustering (Louvain method) was performed using the "FindClusters" function, with a resolution equal to 1.5. Differential expression analysis was performed using the "FindAllMarkers" function to detect markers considering positive log-transformed fold change values above 0.25. Finally, trajectory analysis was performed with the R package "Monocle3" (v.0.2.3.0) (*25*) using CBCs as the source of the pseudotime variable (pseudotime=0). To display trajectory analysis results, UMAP was performed on each matrix using the "reduce_dimension" function found in the Monocle3 package (*26*).

Clusterin lineage tracing

Tamoxifen-inducible *Clu* fate-mapping mice (*Clu*-CreERT; Rosa26-LSL-tdTomato) (*8*) were infected with 150 *Hpb* Larvae. 2.5mg of Tamoxifen (Sigma) was injected intraperitoneally at days 5, 7 and 12 post-infection and small intestines were harvested 14 days post-Tamoxifen injection for histological analysis. tdTomato staining was detected with rabbit anti RFP antibody (1:500; Rockland 600-401-379).

Immune phenotyping

The last 10 cm of the ileum was used to extract lamina propria cells as previously described (*14*). For the Peyer's patches and mesenteric lymph nodes, tissue was crushed in cold HBSS buffer (HBSS supplemented with 2% FBS and 15 mM HEPES), passed through a 100µm filter and centrifuged at 1800rpm for 5 min at 4 °C. Cell suspensions were incubated with a fixable Viability dye (eFluor 506,eBioscience) for 30 min at 4 °C. Cells were then incubated with Fc block (10 min at 4 °C), followed by staining (for 30 min at 4 °C) with the following antibodies in appropriate combinations of fluorophores. From Invitrogen: B220 (RA3–6B2), CD127 (A7R34), CD62L (MEL-14) and GATA-3 (TWAJ). From Biolegend: CD45 (30F11). From BD: CD44 (IM7), CD3 (145-2C11) and CD4 (GK1.5). For staining of intracellular proteins, cells were fixed and permeabilized with the FoxP3 Fix/Perm kit (eBiosciences) according to manufacturer's instructions. Data were acquired with a LSR Fortessa (BD Biosciences) and analyzed using FlowJo software (TreeStar).

IL4 complex (IL4C)

A long-acting form of IL-4 was produced by mixing 10μg recombinant murine IL-4 (Peprotech) with 50μg of neutralizing monoclonal antibody (Clone 11.B11, BioXcell). *Clu* fate-mapping mice were injected with IL4C, intravenously, at days 10, 12 and 14 post *Hpb* infection and with a single 3mg Tamoxifen injection, intraperitoneal, at day 12 post infection. The small intestinal tissue as well as feces for *Hpb* egg counts were harvested at day 12 post Tamoxifen injection (day 24 post *Hpb* infection).

Parasite burden and fecundity

For worm burden assessment, mice were infected with *Hpb* and the small intestine was harvested at day 28 post infection. Adult worms were picked up one by one and enumerated. To measure parasite fecundity, feces were collected from infected mice at day 14 post infection, weighed and placed in 1 mL saturated NaCl solution. The samples were vortexed and left at room temperature for 24h and then placed at 4 °C. The eggs were enumerated and normalized to the volume of the supernatant and weight of feces.

YAP immunohistochemistry

Small intestinal tissues for immunohistochemistry were fixed overnight in formalin at room temperature. Samples were then dehydrated, embedded in paraffin, and sectioned at 4 μM. After antigen retrieval (10 mM Sodium Citrate buffer, pH 6), and blockage of peroxidase activity in 3% H2O2 solution, sections were stained with an active Yap antibody (1:300; Abcam ab205270). Detection of primary antibodies was achieved using the Dako Envision plus system.

Statistical analysis

GraphPad Prism 9 software was used to perform statistical analyses. Student's T test or One-way or two-way Anova were used.

### Supplementary figures

**Fig. S1. Fetal reversion of the intestinal epithelium at the luminal stage of *Hpb* infection. Related to figures 1 and 2.**

(**A**) Epithelial cells were extracted from uninfected or day 14 post-*Hpb* infected small intestines and the expression of several fetal genes was assessed by qPCR. (**B**) RNAscope of *Il33*, *Il1rn* and *Clu* in day 6 and day 14 post-*Hpb* infected small intestinal tissues. (**C**) Heat map of reads per kilobase million (RPKM) was created using the Heatmapper web tool (*15*). (**D**) GSEA of transcripts associated with mature enterocytes, Goblet cells and Paneth cells at day 14 post-*Hpb* infection. N>3; *p<0.05; **p<0.01, ***p<0.005.

**Fig. S2. Fetal- like reprogramming is partially Yap-dependent.**

(**A**) Immunohistochemistry for active YAP, squares mark area in bottom panel, open arrowheads mark nuclear YAP expression; black arrowheads indicate worm presence. (**B-D**) qPCR analysis of *Yap*$^{-/-}$ and littermate control SI organoids stimulated with HES for 24h. (**E-G**) qPCR analysis of *Clu*$^{cre/cre}$and littermate control SI organoids stimulated with HES for 24h. Scale bar: 100μm N>3; *p<0.05, **p<0.01, ***p<0.005.

**Fig. S3. scRNA sequencing of HES-stimulated organoids. Related to figure 3.**

(**A**) UMAP projection plots, pink represents ENR sample, blue represents ENR+HES sample (left panel). Colors represent cells clustered together on the basis of similarity in global gene expression (right panel). (**B**) dot plot of cluster identifying markers (gene expression list can be found in Table S2). (**C-D**) trajectory analysis using CBCs as the source of the pseudotime variable (pseudotime = 0), the black lines represent the branched trajectory, white circle represents the source population, black circles represent branching points and grey circles represent trajectory endpoints.

**Fig. S4. A balanced regulation of the ISC niche by helminths and type 2 immune signaling supports durable infection. Related to figure 4.**

(**A**) representative photomicrographs of *Clu* RNAscope of *Il4ra*$^{WT}$ and *Il4ra*$^{\Delta IEC}$ tissue sections at day 14 post-*Hpb* infection. (**B**) representative photomicrographs of dtTomato IHC for *Clu* lineage tracing in wild type mice treated with PBS of IL4C. (**C**) *Hpb* worm burden at day 28 post-infection in wild type (WT) and *Stat6*$^{-/-}$ mice. (**D**) *Hpb* egg burden following IL4C treatment.

**Fig. S5. Immune phenotyping of *Stat6*$^{-/-}$ and *Il4ra*$^{\Delta IEC}$ at day 14 post-*Hpb* infection.**

Immune phenotyping at day 14 post-*Hpb* infection. (**A**) the small intestine (SI) (**B**) Peyer's patches (PPs) and (**C**) mesenteric lymph nodes (MLN) were harvested from WT and *Stat6*$^{-/-}$ mice. Single cell suspensions were made and flow cytometry performed. Cells were gated for viable CD45+CD3+B220-CD4+CD62L-CD44high cells and GATA3+ were identified as Th2 cells. Percent of cells out of the CD62L-CD44+ population is presented. (**D**) the SI, (**E**) PPs and (**F**) MLN were harvested from *Il4ra*$^{WT}$ and *Il4ra*$^{\Delta IEC}$ mice. Single cell suspensions were made and flow cytometry performed. Cells were gated for viable

CD45+CD3+B220-CD4+CD62L-CD44high cells and GATA3+ were identified as Th2 cells. Percent of cells out of the CD62L-CD44+ population is presented. N=6; *$p<0.05$, **$p<0.01$, ***$p<0.005$.

**Fig. S6. Helminths directly shape the intestinal epithelium to subvert host-derived mechanisms of expulsion.**

During the chronic and luminal stage of *Hpb* infection, adult worms reside in close proximity to the intestinal villi, where they grow, breed and deposit eggs into their host feces. Adult worms secrete a plethora of molecules know as excretory-secretory products (HES) with divergent immune-modulatory functions, especially targeting the type 2 immune response that is responsible for parasitic helminth expulsion. Here we establish that HES can directly target the intestinal stem cell niche to induce a restorative fetal-like reversion of the epithelium, the expansion of *Clu*+ revival stem cells (revSCs) while suppressing the secretory potential (Goblet, Tuft and Paneth cell differentiation) of the stem cell niche. Finally, we show that type 2 cytokines (IL4 and IL13) suppress the HES-induced fetal-like reversion. Collectively, our data suggest that symbiotic parasites such as *Hpb* have evolved with their hosts to co-opt a gut restorative pathway to persist and continue their life cycle. Image created with Biorender.com.

**Table S1.**

Differentially expressed genes ENR vs ENR+HES.xlsx

**Table S2.**

Cluster defining markers.xlsx
